# Severe dysregulation of TNF expression leads to multiple inflammatory diseases and embryonic death

**DOI:** 10.1101/2020.05.28.120501

**Authors:** Elise Clayer, Destiny Dalseno, Andrew Kueh, Derek Lacey, Minhsuang Tsai, Elysa Carr, Verena C. Wimmer, Philippe Bouillet

**Affiliations:** The Walter and Eliza Hall Institute of Medical Research, Parkville, Victoria 3052, Australia; Department of Medical Biology, The University of Melbourne, Melbourne, Victoria 3052, Australia; The Florey Institute of Neuroscience and Mental Health, University of Melbourne, Parkville, Victoria 3052, Australia

**Keywords:** 3’ untranslated region, control of mRNA stability, development, inflammation

## Abstract

Post-transcriptional regulation mechanisms regulate mRNA stability or translational efficiency via ribosomes and recent evidence indicates that it is a major determinant of the accurate levels of cytokine mRNAs. While transcriptional regulation of *Tnf* has been well studied and found to be important for the rapid induction of *Tnf* mRNA and regulation of the acute phase of inflammation, study of its post-transcriptional regulation has been largely limited to the role of the AU-rich element (ARE), and to a lesser extent, that of the constitutive decay element (CDE). We have identified a new regulatory element (NRE) in the 3’ untranslated region (3’UTR) of *Tnf*, and demonstrate that ARE, CDE and NRE cooperate to efficiently down regulate *Tnf* expression and prevent autoimmune inflammatory diseases. We also show for the first time that excessive TNF may lead to embryonic death.

## Introduction

The role of excessive TNF in multiple inflammatory diseases such as rheumatoid arthritis (RA), ankylosing spondylitis (AS) or inflammatory bowel disease (IBD) has been largely documented over the past twenty-five years, and the development of anti-TNF therapies has been a milestone in the treatment of RA and many other inflammatory conditions. Adalimumab, an anti-TNF biologic used to treat RA, has been the best-selling drug worldwide for several years. Many signals have been described that may lead to the increase of TNF expression through the activation of NF-kB. The mechanisms that regulate *Tnf* expression post-transcriptionally, however, have been much less studied, with the notable exception of the action of tristetraprolin (ZFP36/TTP) on the AU-rich element (ARE) found in *Tnf* 3’ untranslated region (3’UTR). Mice with a deletion of the ARE from the *Tnf* gene (TNFDARE) were first described 20 years ago and shown to develop severe arthritis and IBD (1). We recently described BPSM1 mice, a spontaneous mutant mouse in which the insertion of a retrotransposon eliminates most of the 3’UTR of *Tnf*, causing elevated levels of the cytokine and the development of RA and heart valve disease in these mice (2). We also identified a previously unrecognised regulatory element in *Tnf* 3’UTR. We now have further dissected the 3’UTR of *Tnf* and show that several discrete regulatory elements cooperate to efficiently regulate the amount of *Tnf* RNA present in a given cell at a given time. To further evaluate the importance of the 3’UTR in the regulation of TNF levels in vivo, we generated mice with deletions of one or two regulatory elements in *Tnf* untranslated region and show that the phenotypes associated with these deletions vary enormously in severity, one of them even causing embryonic death at E16.5.

## Results

### Three major regulatory elements in *Tnf* 3’UTR

We previously used transient transfection and a series of GFP reporters derived from the pGL3-promoter reporter (Promega) to investigate the potential interaction of members of the CCCH family of zinc finger-containing proteins (ZFP) with several engineered variants of *Tnf* 3’UTR (2). This led to the identification of a new regulatory element located at the distal end of *Tnf* 3’UTR, just before the polyadenylation signal. To further investigate potential interactions between the different regions (regions 1-6, Fig 1A) previously defined, we have generated additional reporter constructs with deletions of multiple regions within *Tnf* 3’UTR and performed transient transfection experiments in HEK293 cells. All constructs were compared to a reporter containing the wildtype 3’UTR of mouse *Tnf* (Fig 1B). Deletion of region 1, 2, 3 or 5 alone had no significant impact on the level of expression of GFP, whereas deletion of region 4 or 6 led to a 3-fold and 2.5-fold increase in GFP expression, respectively. Deletion of regions 1, 2 and 3 together did not lead to a change in reporter activity, suggesting that these three regions do not contain any motif of significance regarding post-transcriptional regulation of *Tnf* expression, at least in the context of these experiments. Deletion of regions 4 and 5 together led to a 5-fold increase in reporter activity, suggesting that regions 4 and 5 somehow cooperate to downregulate *Tnf* expression. Deletion of regions 4 and 6 together led to a 25-fold increase in GFP expression, indicating the crucial role of these two regions in the downregulation of *Tnf* expression. Surprisingly, deletion of regions 4, 5 and 6 together only led to a 10-fold increase in GFP reporter expression, and a construct harbouring the *Tnf* 3’UTR found in the BPSM1 mice (lacking regions 2 to 6 replaced by a retrotransposon) reported a 7.5-fold increase in GFP.

**Figure 1.**
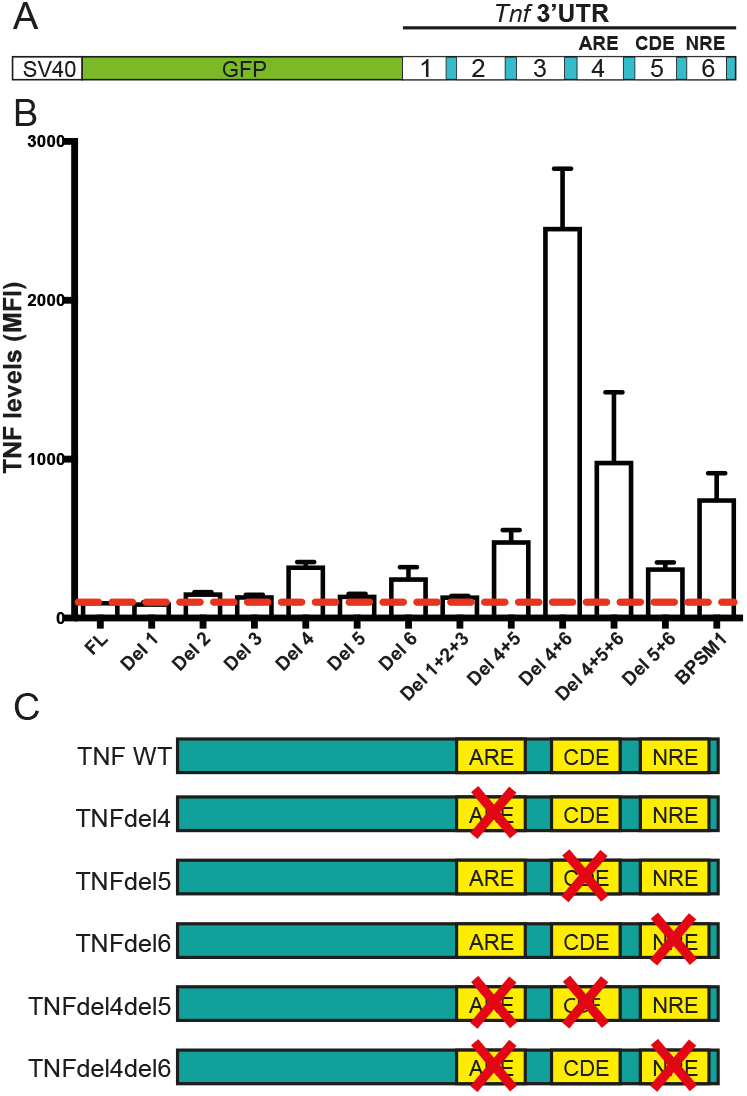
Several elements within *Tnf* 3’UTR cooperate to regulate its expression post-transcriptionally. (A) Schematic representation of the GFP reporter constructs used to assay the effect of the deletion of particular regions of mouse *Tnf* 3’UTR. (B) Deletion of regions 4, 5, 6, or combinations of those, greatly affect the expression of the GFP reporter relative to the full-length (FL) *Tnf* 3’UTR in HEK293 cells. Data represent mean ± SEM of three independent experiments. (C) Schematic representation of the *Tnf* 3’UTR mutations that were engineered in mice using the CRISPR/Cas9 technology.

Together, these results suggest that the most important elements controlling post-transcriptional regulation of *Tnf* expression reside in regions 4, 5 and 6 of the 3’UTR. While all the results concerning regions 4 and 6 suggest that these elements are bound by repressors of *Tnf* expression, the results obtained with the construct lacking regions 4 and 5 and the construct lacking regions 4, 5 and 6 suggest that region 5 might have a dual role in the regulation of *Tnf* expression. Region 5 contains the constitutive decay element (CDE), a 17-nt hairpin loop that is bound by Rc3h1 and Rc3h2 (also known as Roquin 1 and 2) (3). Rc3h1 was described as crucial for the limitation of TNF-a production in resting and activated macrophages (3). Since our region 5 covers more than the CDE itself, it is possible that an as yet undescribed regulatory element lies within the 60 nucleotides of region 5 that are not the CDE. Since our aim was to investigate which regulatory elements are crucial in the regulation of *Tnf* expression, we used these results to guide our choice as to which deletions we would engineer in a series of novel mouse models. We decided on the removal of region 4, region 5 or region 6 alone, and the combined removal of region 4 and 5 or 4 and 6 (Fig 1C).

### TNFdel4 mice

Region 4 of *Tnf* 3’UTR contains the AU-rich elements (ARE) that are bound by TTP to regulate Tnf post-transcriptionally (4). The mutations in TNFDARE and BPSM1 mice are both dominant and lead to RA of similar severity (1, 2). Surprisingly, deletion of region 4 in our reporter system led to a smaller increase in GFP expression than the increase caused by the BPSM1 mutation (Fig 1). Moreover, we have shown that the cloning strategy used to produce TNFDARE mice left in region 6 a 132-nucleotide vector sequence containing a loxP site and several restriction sites. Owing to our results showing a strong cooperation of regions 4 and 6, we suspected that this 132-nt insertion in region 6 might have altered the expression of *Tnf* more than a deletion of the ARE alone would have. Therefore, we generated TNFdel4 mice, in which we deleted the ARE from *Tnf* 3’UTR using CRISPR, carefully leaving any other region of the UTR unmodified. While heterozygous *Tnf^DARE/+^* mice developed arthritis, IBD (1) and heart valve disease (HVD) (2), heterozygous *Tnf^del4/+^* mice developed none of these ailments, even at 18 months of age (Fig2). Homozygous *Tnf^del4/del4^* mice, by contrast, developed oesophagitis, ileitis and colitis (Fig 2), but only mild arthritis, and no heart disease (Fig S1). Arthritis became severe in older *Tnf^del4/del4^* mice (> 200 days-old), but they never showed heart valve problems. While *Tnf^DARE/DARE^* mice succumbed to disease between 5 and 12 weeks of age (1), *Tnf^del4/del4^* mice lived up to one year. Like in TNFDARE mice, concomitant ablation of TNFR1 prevented all pathologies in *Tnf^del4/del4^* mice (data not shown). Thus, it appears that the deletion of the ARE from *Tnf* 3’UTR is less detrimental to mouse health than previously reported, this difference most likely stemming from the insertion of a loxP site and some vector sequences in region 6 of the UTR in TNFDARE mice (see below).

**Figure 2.**
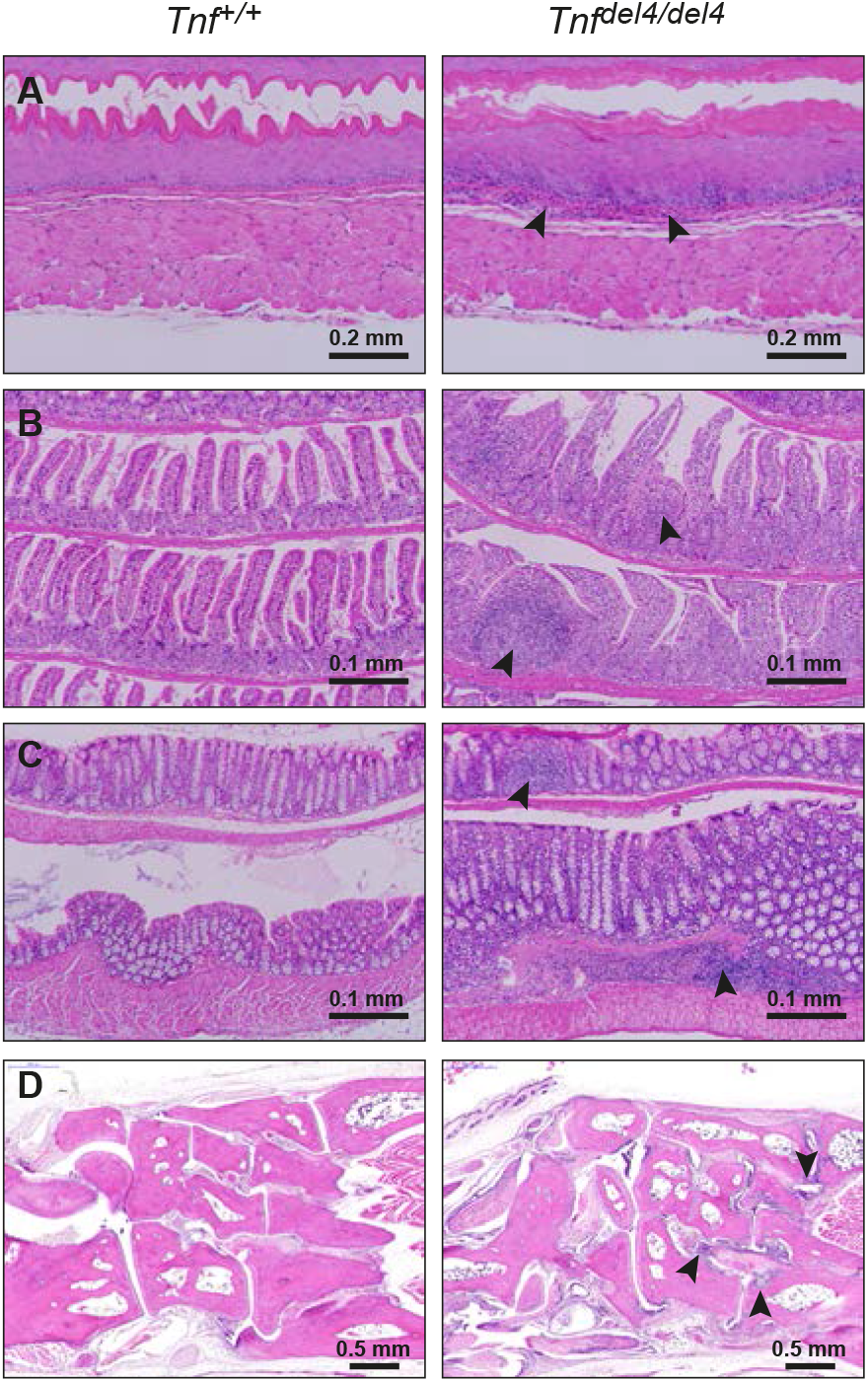
In vivo deletion of the ARE from *Tnf* 3’UTR causes severe IBD, but only mild arthritis. Sections through the oesophagus (A), ileum (B) and colon (C) show the infiltration of numerous immune cells (arrowheads) within these organs in *Tnf^del4/del4^* mice. (D) Sections through the ankle of 200 days-old *Tnf^del4/del4^* mice display signs of moderate arthritis (arrowheads).

We have shown recently that, in addition to RA and heart valve disease, BPSM1 mice (both heterozygotes and homozygotes) also develop tertiary lymphoid organs (TLOs) such as inducible bronchus-associated lymphoid tissue (iBALT) and nodular lymphoid hyperplasia (NLH) in the bone marrow (5). Likewise, both TNFdel4 heterozygous and homozygous mice also developed iBALT and NLH (Fig S1, and data not shown). The absence of any phenotype in *Tnf^del4/+^* mice reinforces our previous conclusion that neither iBALT nor NLH have a pathogenic role, in line with the normal presence of these structures in species such as rats or rabbits.

In addition to these previously described phenotypes, *Tnf^del4/del4^* mice were immediately identifiable even before weaning by their wet/oily-looking fur. This was due to the increase in size of their sebaceous glands (Fig S2), a phenotype that was not present in BPSM1 or TNFDARE mice.

### TNFdel5 mice

Region 5 of *Tnf* 3’UTR contains the constitutive decay element (CDE) (6), a 17-nucleotide stem-loop motif recognized by Rc3h1 (Roquin) and Rc3h2 (Roquin2) (3). While Leppek et al. found that binding of Roquin initiates degradation of *Tnf* mRNA and limits TNF-a production in macrophages, our results with a GFP reporter assay in HEK293 cells suggested that Roquin may have activator function on both mouse and human Tnf 3’UTR, and that Roquin2 may be neutral (on human UTR) or inhibitory (on the mouse UTR) (2). These discrepancies may well be due to the differences between the cell systems used for these experiments. Region 5 also contains a single TATTTAT element that was postulated to be acting as an ARE (7). However, the study focussing on CDE functions suggested that downstream sequences did not affect the translational response of *Tnf* mRNA (6). To assess the role of the CDE on *Tnf* expression in vivo, we deleted region 5 of *Tnf* 3’UTR by CRISPR to generate the TNFdel5 strain. *Tnf^del5/+^* and *Tnf^del5/del5^* mice failed to develop any obvious inflammatory phenotype in an 18-months observation period (data not shown), suggesting that the CDE by itself only has a minor role in regulating the steady state expression of *Tnf*.

### TNFdel6 mice

We have shown that the 76 nt-long region 6 of *Tnf* 3’UTR contains a new regulatory element (NRE) that strongly cooperates with the ARE in the downregulation of *Tnf* expression. In our GFP reporter system in HEK293 cells, region 6 appeared involved in the downregulation of *Tnf* by the RNA binding proteins Zc3h12a and Zc3h12c (2). To narrow down this effect to a shorter motif within region 6, we have generated six additional reporter constructs missing both region 4 and a smaller part of region 6. None of these constructs showed a synergy similar to that observed with the construct missing region 4 and the entire region 6 (data not shown). It is however possible that region 6 simply participates in the overall secondary structure of *Tnf* 3’UTR. We used CRISPR to delete the 76-nucleotide region 6 from *Tnf* 3’UTR and generate the TNFdel6 strain. *Tnf^del6/+^* and *Tnf^del6/del6^* mice also failed to develop any obvious inflammatory phenotype within 2 years (data not shown), demonstrating that deleting the NRE alone is not sufficient to alter *Tnf* expression enough to cause an inflammatory disease.

### TNFdel4del5 mice

The deletion of both regions 4 and 5 from *Tnf* 3’UTR resulted in higher GFP expression than the deletion of either region alone in our reporter system (Fig 1). We thus used CRISPR to generate the TNFdel4del5 strain with the *Tnf* 3’UTR lacking both regions. *Tnf^del4del5/+^* mice presented at weaning with a wet/oily-looking fur (Fig S2) and a slightly smaller size than their WT littermates, a phenotype similar to that observed in *Tnf^del4/del4^* mice. *Tnf^del4del5/+^* mice developed oesophagitis, ileitis and colitis, as well as iBALT and NLH (Fig 3, Fig S3, and data not shown). Unlike *Tnf^del4/del4^* mice however, they also developed arthritis and heart valve disease at an early age (Fig S3), and rarely reached 250 days of age. Thus, the concomitant deletion of both regions 4 and 5 from *Tnf* 3’UTR drastically increases the severity of the consequences due to the deletion of region 4 alone and shows that regions 4 and 5 of *Tnf* 3’UTR cooperate *in vivo* to regulate the expression of *Tnf*. The complete absence of the heart valve phenotype in TNFdel4 mice and its presence in TNFdel4del5 mice suggests that the regulation also involves cell type specificity, maybe due to tissue-specific expression of the proteins that recognize these different regulatory elements of *Tnf* 3’UTR.

**Figure 3.**
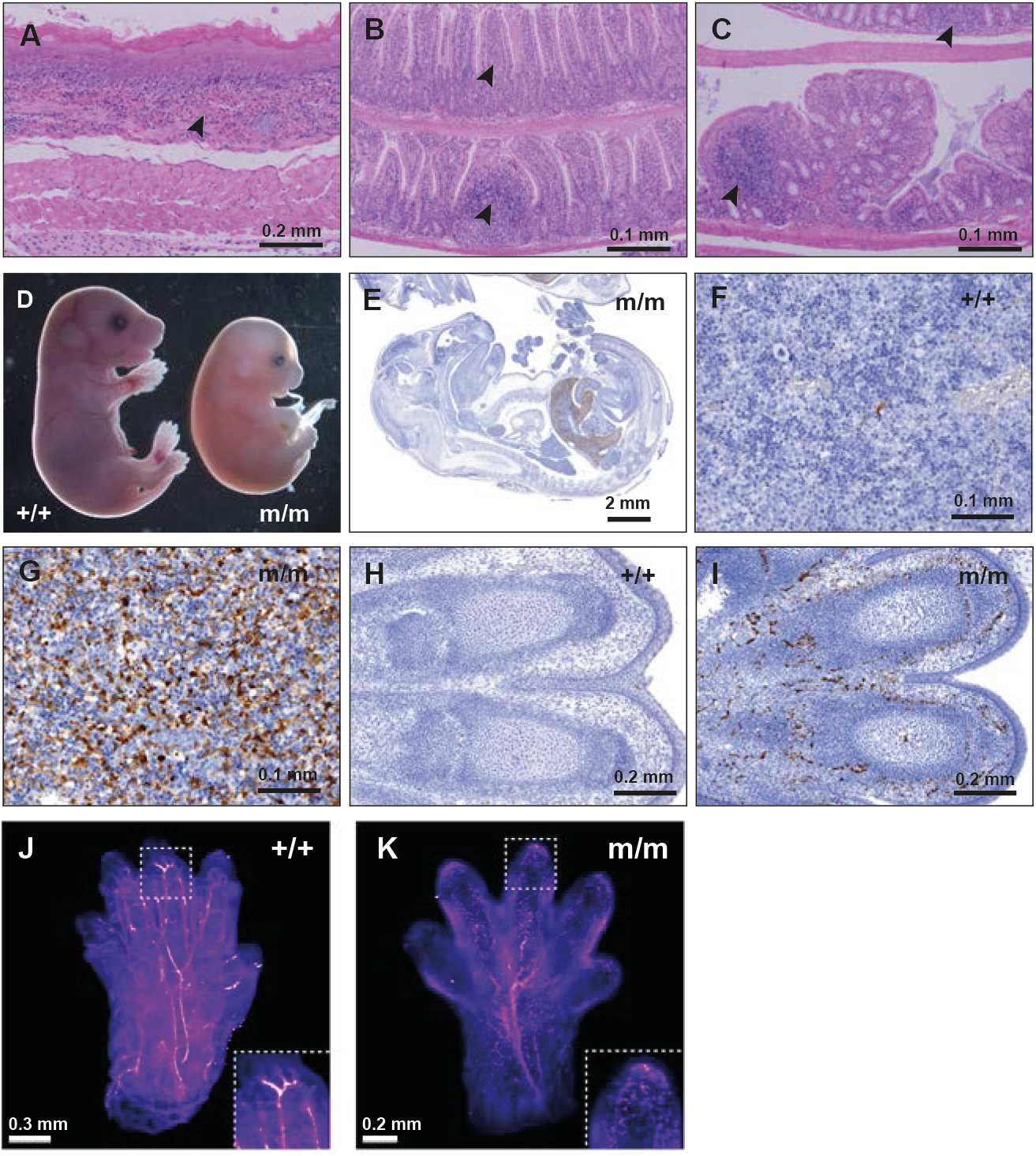
Concomitant deletion of region 5 increases the severity of the inflammatory phenotype due to the removal of the ARE from *Tnf* 3’UTR. Oesophagitis (A), ileitis (B) and colitis (C) in a 100 days-old *Tnf^del4del5/+^* mouse. Arrowheads indicate immune cell infiltration in these tissues. (D) Homozygote *Tnf^del4del5/del4del5^* (m/m) embryos die at E16.5 and appear devoid of major blood vessels. (E) Active Caspase-3 staining of an E15.5 homozygote TNFdel4del5 embryo (m/m) shows extensive cell death in the liver. Active Caspase-3 stained liver of WT (F) and homozygote (G) E16.5 TNFdel4del5 embryos at 40X magnification. 20X magnification shows death of blood vessels in the digits of an E16.5 TNFdel4del5 homozygote embryo (I) and the absence of cell death in WT embryo (H) at the same stage of development. (J, K) Label free lightsheet imaging of hemoglobin autofluorescence in cleared front paws of WT (+/+) and homozygote *Tnf^del4del5/del4del5^* (m/m) E16.5 embryos. Still image from Supplementary video 1.

No *Tnf^del4del5/del4del5^* homozygote mouse was detected in the litters from heterozygote parents. Therefore, we arranged timed matings to determine the time of death of homozygous embryos. Examination of these embryos indicated that development proceeded normally until E15.5. At E16.5 however, homozygous embryos appeared smaller, and their limbs, heads and tails were very pale due to the disappearance of major blood vessels that were clearly visible in their WT and heterozygous littermates (Fig 3). No homozygous embryo was found alive after E16.5. Light sheet microscopy clearly showed the fragmentation of the vascular system in the limbs of these embryos (Supplementary video 1). Staining histological sections for active caspase 3 confirmed the death of endothelial cells in the limbs and revealed massive apoptosis of hepatocytes in the liver, which was also devoid of most blood vessels (Fig 3). In addition, the proportion of nucleated erythroblasts was considerably higher in *Tnf^del4del5/del4del5^* homozygote embryos than in controls (Fig S3). We conclude that *Tnf^del4del5/del4del5^* homozygote embryos die of vascular collapse and liver degeneration around E16.5.

Loss of TNFR1 once again completely prevented all the pathologies observed in *Tnf^del4del5/+^* and *Tnf^del4del5/del4del5^* animals, and compound *Tnf^del4del5/del4del5^*/*TNFR1*^-/-^ mice lived and reproduced without developing any symptoms (data not shown). Interestingly, loss of a single allele of *TNFR1* also prevented the embryonic death of *Tnf^del4del5/del4del5^* animals. However, *Tnf^del4del5/del4del5^/TNFR1*^+/-^ mice developed arthritis, heart valve disease and IBD (Fig S4), and only lived to ~100 days. This result exemplifies the delicate balance between TNF and TNFR1 in the signalling of this cytokine.

Altogether, these data show that, as suggested by the results in our GFP reporter system, regions 4 and 5 of *Tnf* 3’UTR cooperate to regulate expression of TNF *in vivo* and that this cooperation is necessary to prevent embryonic death.

### TNFdel4del6 mice

Deletion of regions 4 and 6 from *Tnf* 3’UTR led to the highest level of GFP expression in our reporter system. Therefore, we expected that the phenotype associated with this double deletion *in vivo* would be remarkable. To generate TNFdel4del6 mice, we first attempted to create the additional TNFdel6 mutation in *Tnf^del4/del4^* embryos, or to create the additional TNFdel4 mutation in *Tnf^del6/del6^* embryos. These approaches failed to generate viable animals harbouring both mutations on the same allele, suggesting that the TNFdel4del6 mutation may be lethal on a TNFR1 WT background. We then designed a new strategy which involved generating the double deletion simultaneously with a repair template and used this approach in WT embryos and in *Tnfr1^-/-^* embryos. While no animal bearing the double mutation was generated on the WT background, *Tnf^del4del6/del4del6^/Tnfr1^-/-^* and *Tnf^del4del6/+^/Tnfr1^-/-^* animals were generated successfully and did not present with any phenotype (data not shown). To examine the effects of the homozygous deletion of both regions 4 and 6 *in vivo*, we crossed *Tnf^del4del6/+^/Tnfr1^-/-^* animals with WT partners and examined their progeny. About 80% of *Tnf^del4del6/+^/Tnfr1^+/-^* animals died within a few hours after being born. Histopathological analysis indicated that these animals had failed to initiate respiration, and all showed underinflated lungs (Fig 4). Twelve *Tnf^del4del6/+^/Tnfr1*^+/-^ animals survived to 20 days. These animals were very runty and showed signs of arthritis (deformed wrists and ankles) from 7 days of age, but their fur did not look wet or oily. Histological examination of 20-day old *Tnf^del4del6/+^/Tnfr1*^+/-^ animals revealed the presence of IBD in the gut, large pannus tissue invasion in the synovial joints, iBALT in the lungs, but only limited inflammation was seen in the heart valves or aortic root (Fig 4, and data not shown).

**Figure 4.**
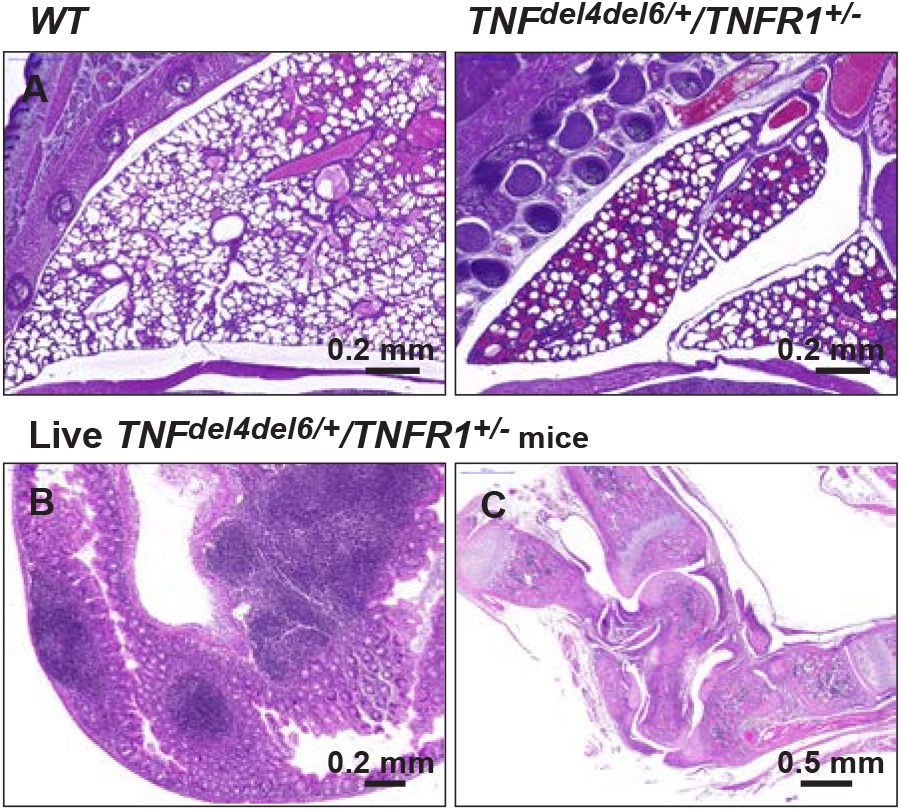
Phenotypes due to the combined deletion of regions 4 and 6. (A) Underinflation of lungs is the cause of the death of most *Tnf^del4del6/+^/Tnfr1*^+/-^ pups soon after birth. *Tnf^del4del6/+^/Tnfr1*^+/-^ pups that survive develop IBD (B) and extremely severe arthritis (C). (B) Immune cell infiltration in the caecum of a 20 days-old *Tnf^del4del6/+^/Tnfr1*^+/-^ pup as seen by H&E staining. (C) Dislocated and pannus-infiltrated ankle in the same animal as in (B).

The severity of the phenotype in *Tnf^del4del6/+^/Tnfr1*^+/-^ animals prevented us from generating *Tnf^del4del6/del4del6^/Tnfr1*^+/-^ animals or *Tnf^del4del6/+^/Tnfr1*^+/+^ animals.

Even though the deletion of region 6 had no consequence on the general health, the concomitant deletion of regions 4 and 6 of *Tnf* 3’UTR *in vivo* led to the most severe phenotype since it was observed in heterozygous mice with a single functional *Tnfr1* allele.

To estimate how these mutations affected the levels of circulating TNF in vivo, we took advantage of the fact that the concomitant loss of Tnfr1 prevented all the pathologies due to an increase in TNF expression. On the *Tnfr1*^-/-^ background, we were able to generate healthy adult mice with heterozygote or homozygote TNFdel4del5 or TNFdel4del6 mutations. The ELISA results (Fig 5) show that the levels of TNF in the serum are strikingly similar to those predicted by the reporter assays in HEK293 cells described in Figure 1. Levels of circulating TNF however are not sufficient to explain some of our observations. For example, the levels of TNF in homozygous BPSM1 (m/m) and TNFdel4del6 (both m/+ and m/m) mice are superior to those of TNFdel4del5 homozygous mutants, yet neither BPSM1^m/m^ nor TNFdel4del6 (m/+ at least) mutant mice die at E16.5 as TNFdel4del5^m/m^ do.

**Figure 5.**
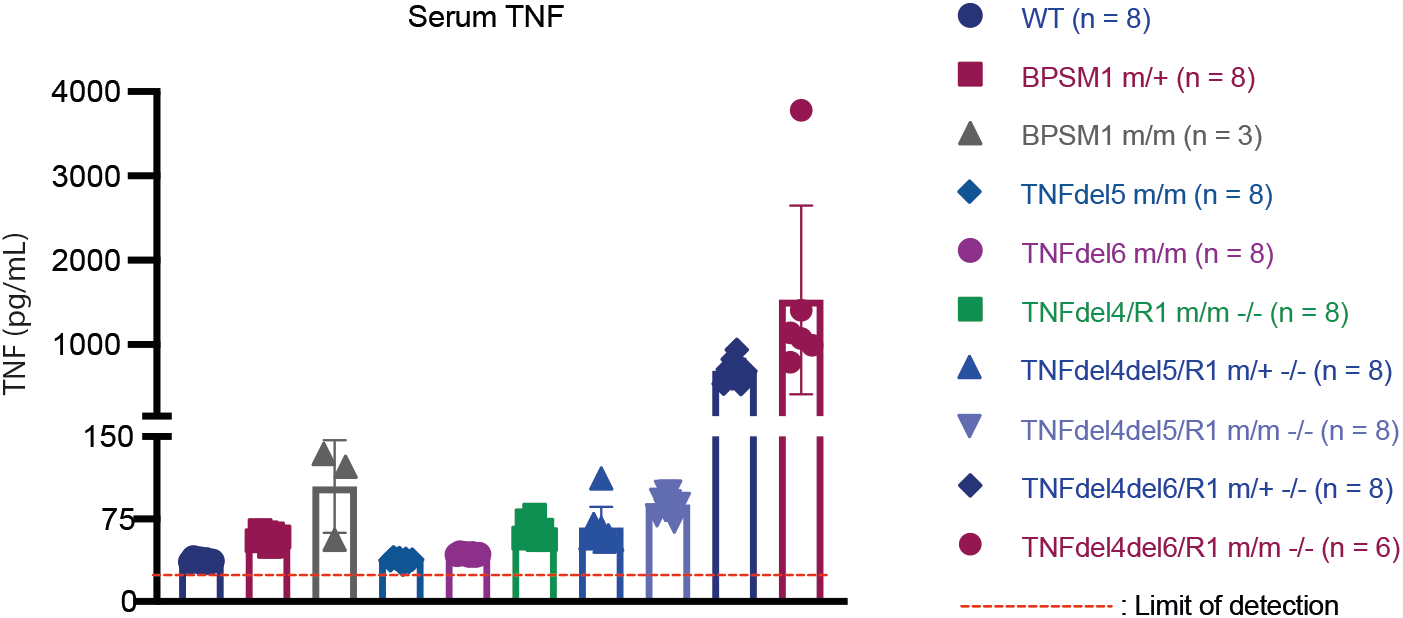
Serum TNF levels vary greatly according to mutations in *Tnf* 3’UTR. TNF levels were determined by ELISA as described. Data represent mean ± SEM of the number of animals indicated.

### Hematopoietic reconstitution

We have shown previously that the hematopoietic system of lethally irradiated wildtype C57BL/6 recipients can be successfully reconstituted with bone marrow cells from *BPSM1^m/m^/TNFR1*^-/-^ healthy animals, and that the recipient mice proceed to develop the disease within two months following transplantation (2). We performed similar experiments with *TNF^del4/del4^/TNFR1*^-/-^, *TNF^del4del5/del4del5^/TNFR1*^-/-^ and *TNF^del4del6/del4del6^/TNFR1*^-/-^ as healthy bone marrow donors. Recipients of *TNF^del4/del4^/ TNFR1*^-/-^ bone marrow cells had developed NLH, mild HVD, mild IBD and mild arthritis five months after transplantation (Fig S5). Recipients of *TNF^del4del5/del4del5^/TNFR1*^-/-^ cells reconstituted their hematopoietic system, but they became sick three weeks after transplantation. Histological analysis showed that they had developed severe IBD, particularly colitis (Fig S5). Recipients of *TNF^del4del6/del4del6^/TNFR1*^-/-^ cells became ill 7-9 days following transplantation, with clear signs of anaemia and bone marrow failure (Fig S5). Histological examination revealed the occurrence of acute liver necrosis in all mice transplanted with *TNF^del4del6/del4del6^/TNFR1*^-/-^ cells (Fig S6). Thus, it appears that the severity of the phenotypes developing after hematopoietic reconstitution largely depends on the amount of TNF produced by donor cells and the tolerance of certain organs (e.g. liver, gut in particular) to high levels of TNF.

## Discussion

Adenylate-uridylate-rich elements (AU-rich elements; AREs) found in the 3’UTR of many messenger RNAs have long been recognized as the most common determinants of RNA stability in mammalian cells (8, 9). A number of RBPs bind to AREs and stabilize the transcripts while others facilitate their degradation (10). Recent technological advances have allowed the identification of more than 1500 RBPs, which are involved in the maturation, stability, transport and degradation of cellular RNAs (11). RBPs may directly bind to RNA or be integral parts of macromolecular protein complexes that bind to RNA, such as the RNA exosome. Despite the growing amount of data collected on RBPs, many questions remain unanswered. Our understanding of how binding specificity is achieved and how the regulatory function of an individual RBP is influenced by synergy and competition with other RBPs is still far from resolved. In addition, more than 100 different types of RNA modifications have been identified, mostly in the form of naturally occurring modified nucleosides (12). While most of these modifications were discovered in abundant molecules such as tRNAs, rRNAs and small non coding RNAs, recent studies have shown that they occur in all cellular RNAs, including mRNAs, and that they can be reversible (13). The challenge of the new field of epitranscriptomics is to understand the functional consequences of mRNA modifications (14).

The ARE present in *Tnf* 3’UTR has been identified as a crucial part of the post-transcriptional regulation of this gene (15), and a first mouse model, TNFDARE, demonstrated the importance of this type of regulation for the prevention of auto-inflammatory diseases such as RA and IBD (1). Tristetraprolin (TTP) was found to bind *Tnf* ARE, and its ablation from the mouse genome also led to autoinflammatory symptoms, partly due to *Tnf* dysregulation (4).

The discovery that the insertion of a retrotransposon in *Tnf* 3’UTR was the cause for the development of chronic polyarthritis and heart valve disease in BPSM1 mice (2) prompted us to reconsider the role of the 3’UTR in the regulation of TNF levels. Three regions of *Tnf* 3’UTR seem to act in concert to regulate *Tnf* expression post-transcriptionally. The genetic models described here provide strong evidence that post-transcriptional regulation of TNF expression is a critical mechanism to maintain low levels of serum TNF in mice. The variety and the severity of the phenotypes highlight the importance of post-transcriptional regulation in the control of TNF expression in health and disease. In addition, this is the first demonstration that excessive TNF can cause embryonic or perinatal death.

Many studies have documented the role of numerous transcription factors such as the nuclear factor of activated T cells (NFAT) and NF-kB on *Tnf* promoter to explain the sudden increase of TNF in response to many stimuli (16). Early studies of human TNF-transgenic mice showed that, irrespective of the promoter used, the replacement of *TNF* 3’UTR in the transgene with that of the b-globin mRNA was sufficient to drive *TNF* promoter-driven TNF overexpression and the development of tissue-specific inflammation in mice (17). TNF promoter was shown to be constitutively active in a number of cell lines (Hela, NIH3T3 and L929) and that *TNF*3’UTR effectively cancelled reporter gene expression in Hela and NIH3t3 cells, but not in L929 cells (18). In HEK293 cells, we have shown that *Tnf* promoter is able to drive a robust expression of GFP, and that the expression of GFP can be extinguished by swapping the SV40 polyadenylation sequence for the 3’UTR of *Tnf* in the reporter (Fig S7). Although this certainly does not exclude a role for transcription factors in boosting TNF expression at the promoter level in inflammatory conditions, it also opens the possibility that some of these transcription factors could produce a similar result by inhibiting the expression of some of the RBPs that participate in the down-regulation of TNF levels through the 3’UTR of its mRNA.

The increase in the size of sebaceous glands and the associated oily-looking fur in TNFdel4 and TNFdel4del5 mice is rather surprising since it was not observed in TNFDARE, BPSM1 and TNFdel4del6 mice. A recent report indicated that sebaceous gland homeostasis was regulated by a population of innate lymphoid cells (ILCs) residing within hair follicles in close proximity to the glands (19). These RORgt+ ILCs were found to negatively regulate sebaceous gland function by expressing TNF and lymphotoxins LTa3 and LTa1b2, which downregulated Notch signaling. The authors reported that sebaceous glands were hyperplastic in the triple knockout (*Tnf*^-/-^, *Lta*^-/-^, *Ltb*^-/-^) mice, while none of the single knockout showed the phenotype. The fact that TNF overexpression can generate a similar phenotype certainly warrants further investigation.

Our results provide strong evidence that several RBPs participate in the post-transcriptional regulation of *Tnf* mRNA levels in vivo, and that the differential distribution of these proteins within tissues and cell types is probably the cause of the phenotypic variations observed in our mutant mice. The disintegration of the vascular system in TNFdel4del5 homozygote E16.5 embryos suggests that a particular RBP is present in these cells to prevent an excessive expression of TNF, and that this RBP acts through regions 4 and/or 5 of *Tnf* 3’UTR. It is remarkable that none of the other mutant mice, in particular those that express higher levels of circulating TNF, shows this phenotype. Whether these cell type-specific RBPs act as part of a ribonuclease complex such as the RNA exosome (20) or the CCR4-NOT complex (21) will of course require their identification. Interfering with the TNF post-transcriptional regulatory system may represent a novel therapeutic avenue for manipulation of TNF expression. Indeed, Allotrap 1258, a synthetic peptide derived for from the heavy chain of HLA Class I, inhibited concanavalin A- and LPS-induced human and mouse TNF production *in vitro* and *in vivo* (22). Allotrap 1258-mediated inhibition required the presence of *Tnf* 3’UTR to be effective, suggesting that it may activate a component of the RNA degradation complex for its activity. The clear parallels between our *in vivo* results and the results obtained with our reporter system in HEK293 cells indicate that these cells contain at least the critical components of the *Tnf* post-transcriptional regulatory system, suggesting that our reporter system could be used as a basis for a screening strategy aimed at identifying novel regulators of TNF expression.

## Materials and Methods

### Mice

All animal experiments were conducted with the approval of the Animal Ethics Committee of the Walter and Eliza Hall Institute. BPSM1 mice were the result of a spontaneous mutation. Deletions within *Tnf* 3’UTR were engineered with CRISPR/Cas9 technology and verified by sequencing. Conclusions were reached after examining at least ten mice of each genotype. All mice were on the C57BL/6 genetic background.

### TNF 3’UTR reporter assays

GFP reporter constructs engineered as in (Lacey et al. 2015) contained an SV40 early promoter driving the expression of eGFP. Murine *Tnf* 3’UTR (WT, BPSM1-derived, or containing deletions of regions 1-6 as indicated) were inserted between the Xba1 and BamH1 sites (complete sequences upon request). HEK293 cells were transiently transfected using Fugene 6 (Promega) with GFP-*Tnf* 3’UTR reporter constructs and a pGL3-mCherry control construct, and analyzed 3 days later using flow cytometry on a LSR IIW (BD Biosciences). GFP Mean fluorescence intensity was calculated on live mCherry-positive cells and compared to empty vector control.

### TNF ELISA

Secreted TNF was measured from mouse serum using the TNF alpha mouse uncoated ELISA kit (ThermoFisherScientific) and read using the Chameleon plate reader (Hidex, Turku, Finland).

### Bone marrow transplantation experiments

Wildtype C57BL/6 recipients were lethally irradiated (2x 550 rad) and injected with 1-2 x 10^6^ BM cells.

### Tissue clearing and lightsheet microscopy

E15.5 embryo paws were cleared using passive clarity (PACT; (23)). In brief, embryos were harvested and fixed overnight in 4% paraformaldehyde in PBS, pH 7.4. The paws were resected and immersed in PACT monomer solution containing 4% acrylamide (Biorad) and Polymerization Thermal Initiator VA044 (0.25% w/v final concentration; Wako) in PBS overnight at 4dC. The gel was set for 3hrs at 37°C and the tissue was moved into borate-buffered clearing solution (8% (wt/vol) SDS and 50mM sodium sulphite in 0.2M boric acid buffer, pH 8.5). Once cleared, the tissue was washed by repeatedly diluting the clearing solution (1:1) with borate-buffered wash solution (1% TritonX-100 SDS and 50mM sodium sulphite in 0.2M boric acid buffer, pH 8.5) to gradually remove the SDS. The paws were mounted in 1% low melting point agarose and immersed in EasyIndex solution (LifeCanvas) for refractive index matching.

The tissue was imaged using a Zeiss Z.1 lightsheet microscope equipped with a 5x/0.16 detection objective. Blood vessels were detected using hemoglobin autofluorescence excited with the 405nm laser line and detected using a 500-545nm GFP band pass filter (24). Multiview data sets were acquired at 120-degree angles and fused using the Multiview Reconstruction plugin in FIJI (25). 3D reconstruction was performed in Imaris (Bitplane).

## Abbreviations

TNF: tumor necrosis factor
RA: rheumatoid arthritis
IBD: inflammatory bowel disease
3’UTR: 3’ untranslated region
BMDM: bone marrow-derived macrophage
ARE: AU-rich element
CDE: constitutive decay element
HVD: heart valve disease
NLH: nodular lymphoid hyperplasia
iBALT: inducible bronchus-associated lymphoid tissue
GFP: green fluorescent protein

## Acknowledgements

We thank J. Stanley, T. Kitson. M. Watters, and G. Siciliano for animal care and expertise; J. Corbin, K. Weston and T. Nikolaou for automated blood analysis; T. Mak (The Campbell Family Institute for Breast Cancer Research) for TNFR1 KO mice; J. Silke for TNF KO mice. This work was supported by the Australian NHMRC (Program Grant 461221, Research Fellowship 1042629, Project grant 1127885), the Leukemia and Lymphoma Society (Specialised Center of Research grant 7015), the Arthritis Australia Zimmer fellowship, and infrastructure support from the NHMRC (IRISS) and the Victorian State Government (OIS). The generation of TNFdel4, TNFdel5, TNFdel6, TNFdel4del5 and TNFdel4del6 mice used in this study was supported by the Australian Phenomics Network (APN) and the Australian Government through the National Collaborative Research Infrastructure Strategy (NCRIS) program.

---

**Figure S1.**
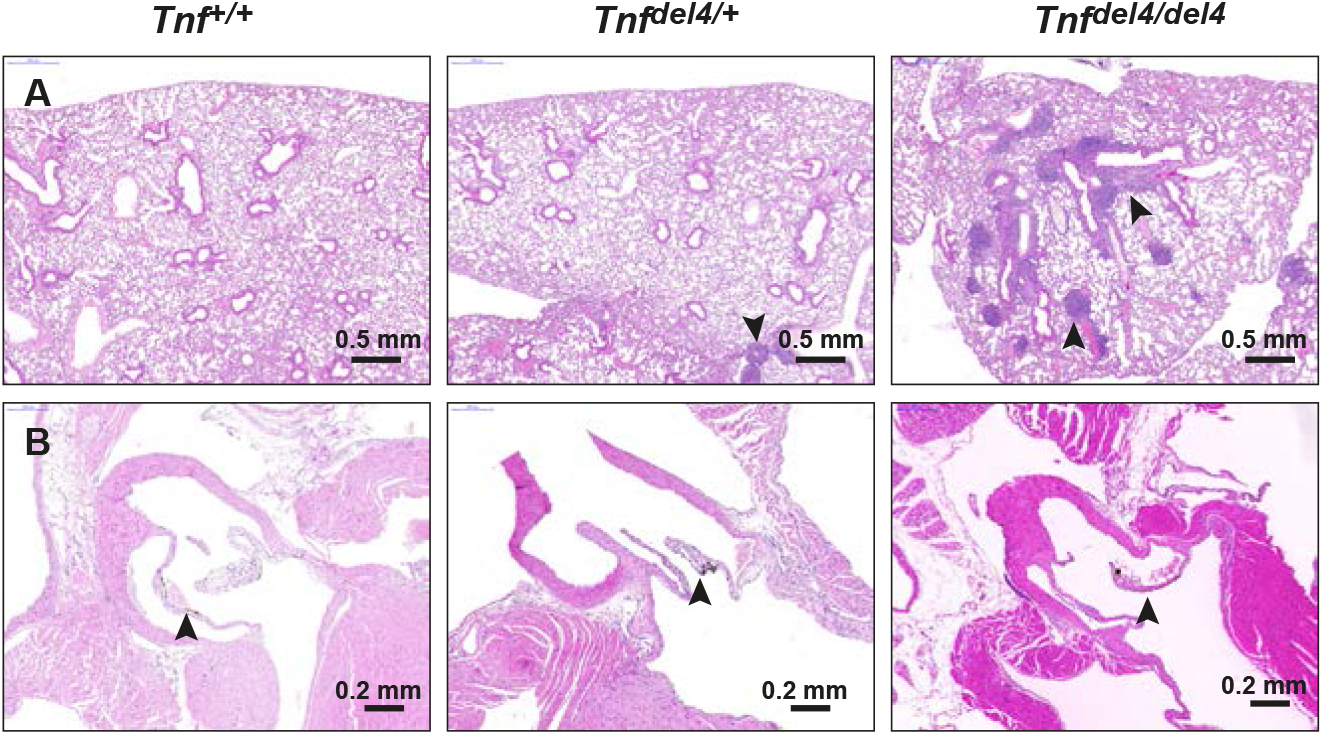
Phenotype of TNFdel4 mice. (A) H&E sections showing the presence of iBALT in the lungs of *Tnf^del4/+^* and *TNF^del4/del4^* mice. (B) H&E sections showing the absence of heart valve disease in 200 days-old *Tnf^del4/+^* and *TNF^del4/del4^* mice.

**Figure S2.**
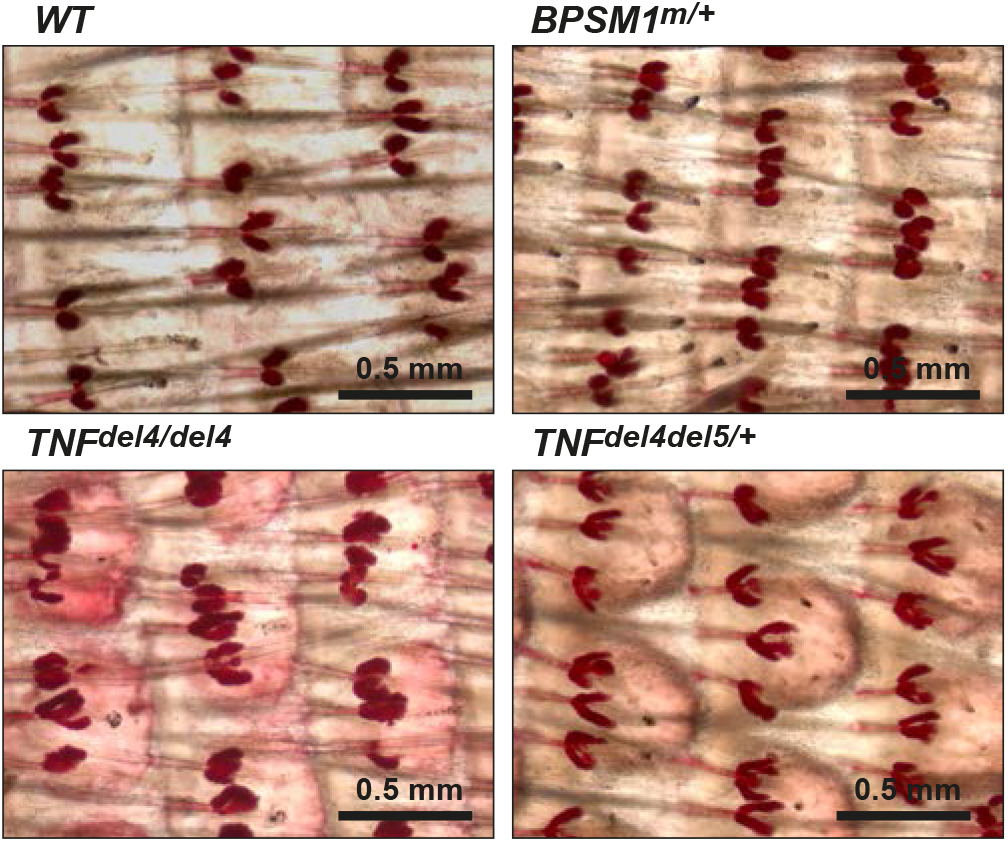
Increased size of sebaceous glands in *Tnf^del4/del4^* and *Tnf^del4del5/+^* mice. Oil red O staining of the tail skin shows the increased size and complexity of the sebaceous glands in *Tnf^del4/del4^* and *Tnf^del4del5/+^* mice.

**Figure S3.**
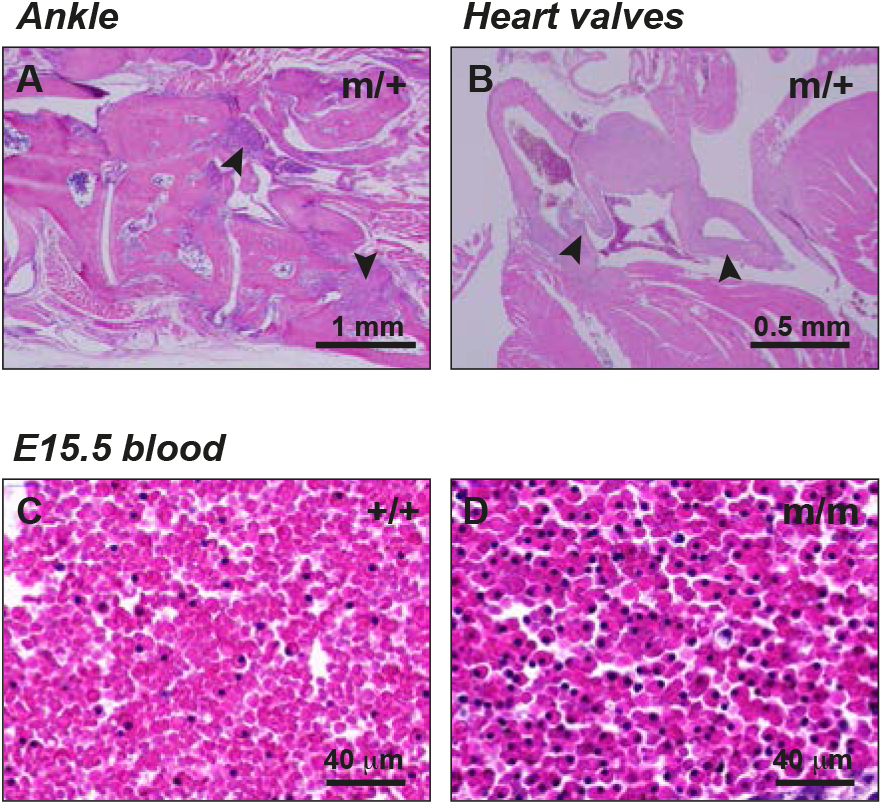
Additional phenotypes in TNFdel4del5 mice. H&E sections arthritis (A) and heart valve disease (B). (C, D) H&E staining of blood of E15.5 embryos shows a large proportion of nucleated erythrocytes in homozygote TNFdel4del5 embryo (m/m, D) compared to a WT embryo at the same stage of development (+/+, C).

**Figure S4.**
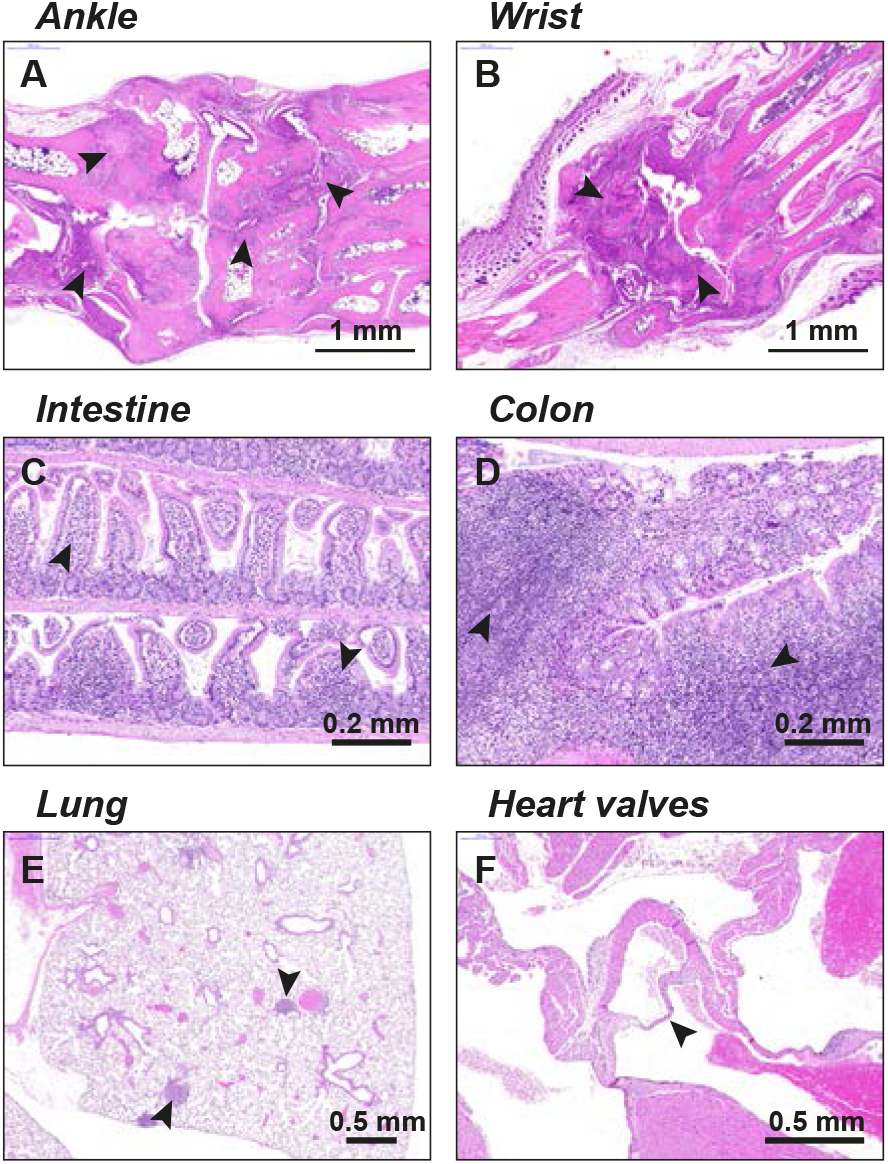
Loss of one TNFR1 allele prevents the embryonic death of *Tnf^del4del5/del4del5^* mice. 100 days-old *Tnf^del4del5/del4del5^*/*TNFR1*^+/-^ animals prevent with very severe arthritis in the ankles (A) and wrists (B), severe IBD (C, D), as well as iBALT in the lungs (E). Arrowheads indicate pannus tissue in (A) and (B), immune cell infiltration in (C) and (D), and iBALT in (E). 100 days-old *Tnf^del4del5/del4del5^*/*TNFR1*^+/-^ animals do not develop heart valve (arrowhead) disease (F).

**Figure S5.**
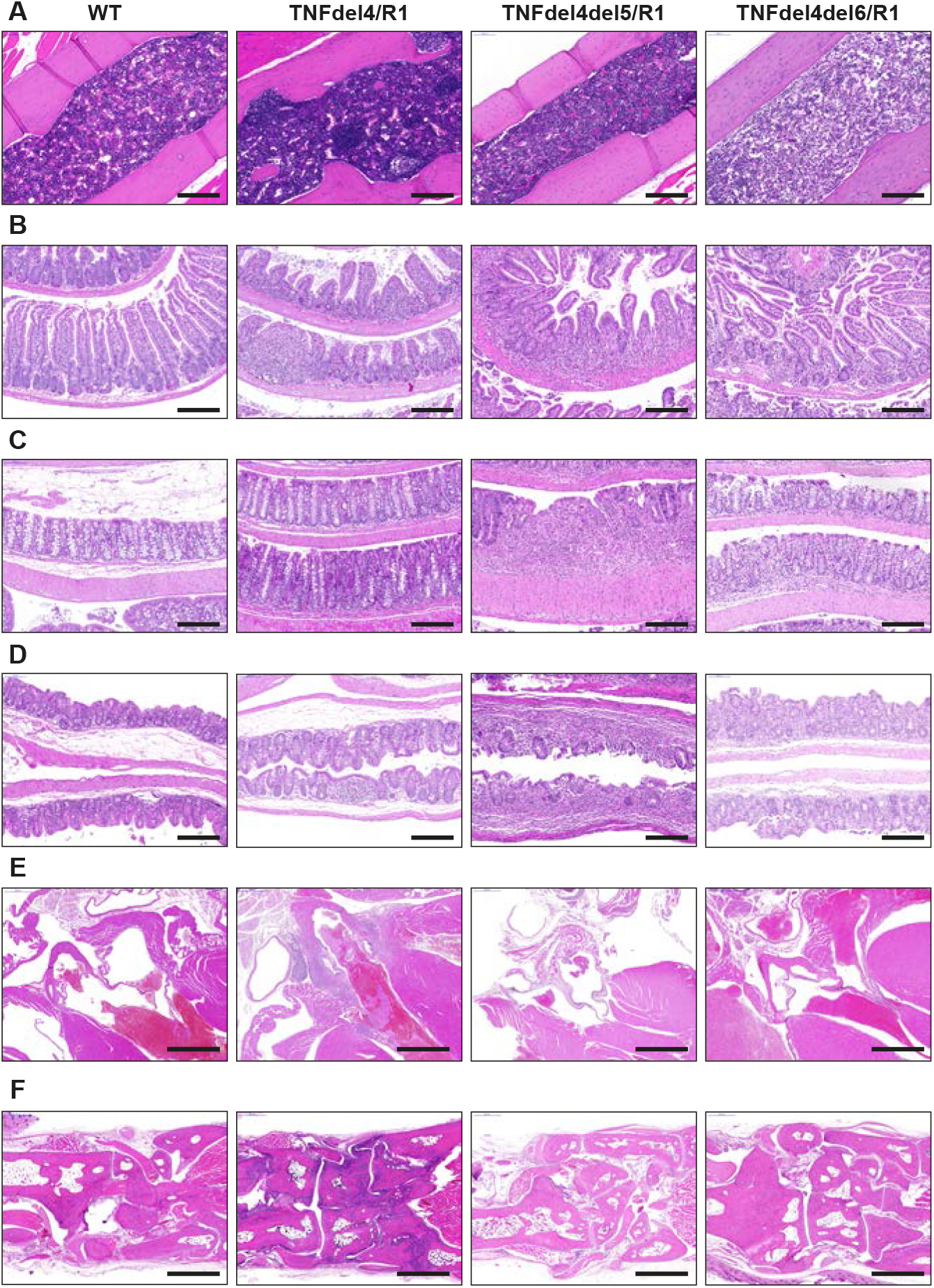
Hematopoietic reconstitution. H&E-stained sections through bone (A), intestine (B), colon (C), caecum (D), heart valves (E) and ankle (F) of lethally irradiated WT C57BL/6 recipient mice transplanted with 1-2 10^6^ bone marrow cells from WT, *Tnf^del4/del4^/TNFR1*^-/-^ (TNFdel4/R1), *Tnf^del4del5/del4del5^/TNFR1*^-/-^ (TNFdel4del5/R1) or *Tnf^del4del6/del4del6^/TNFR1^-/-^* (TNFdel4del6/R1) donors. Recipients of WT and TNFdel4/R1 BM cells were examined 5 months post transplantation. Recipients of TNFdel4del5/R1 BM cells became sick and were examined 21 days post transplantation. Recipients of TNFdel4del6/R1 BM cells failed to reconstitute their hematopoietic system and were examined 9 days post transplantation. Bars: 0.2mm in A, B, C and D; 0.5 mm in E and F.

**Figure S6.**
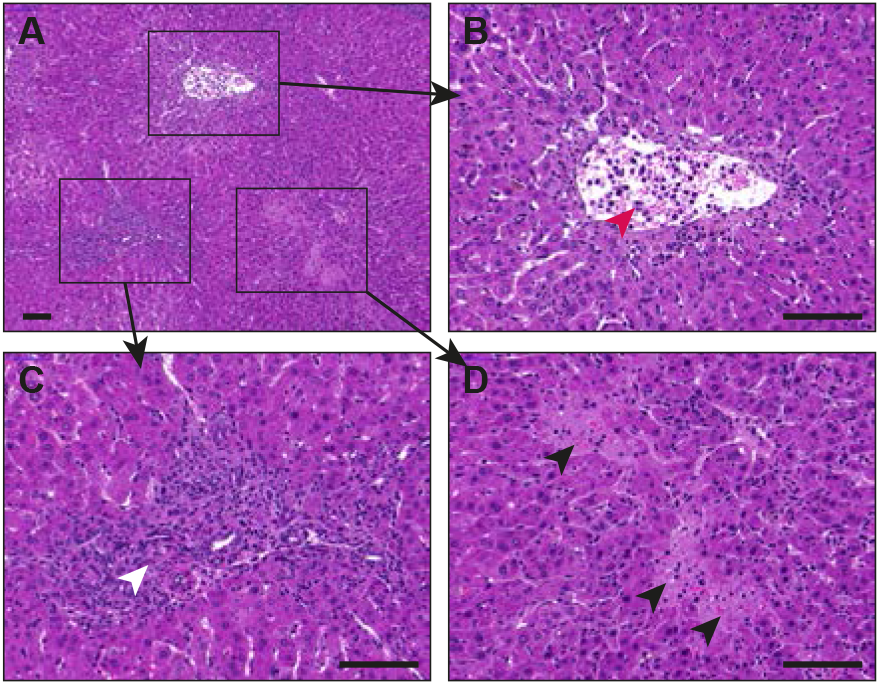
Acute liver necrosis in the recipients of TNFdel4del6/R1 BM cells. (A) H&E-stained section of the liver of a lethally-irradiated WT recipient of TNFdel4del6/R1 BM cells shows the presence of blood vessels filled with the remnants of dead hepatocytes (B, red arrowhead), perivascular infiltrates of immune cells (C, white arrowhead) and necrotic zones devoid of live cells (D, white arrowheads). Bars: 100μm in all panels.

**Figure S7.**
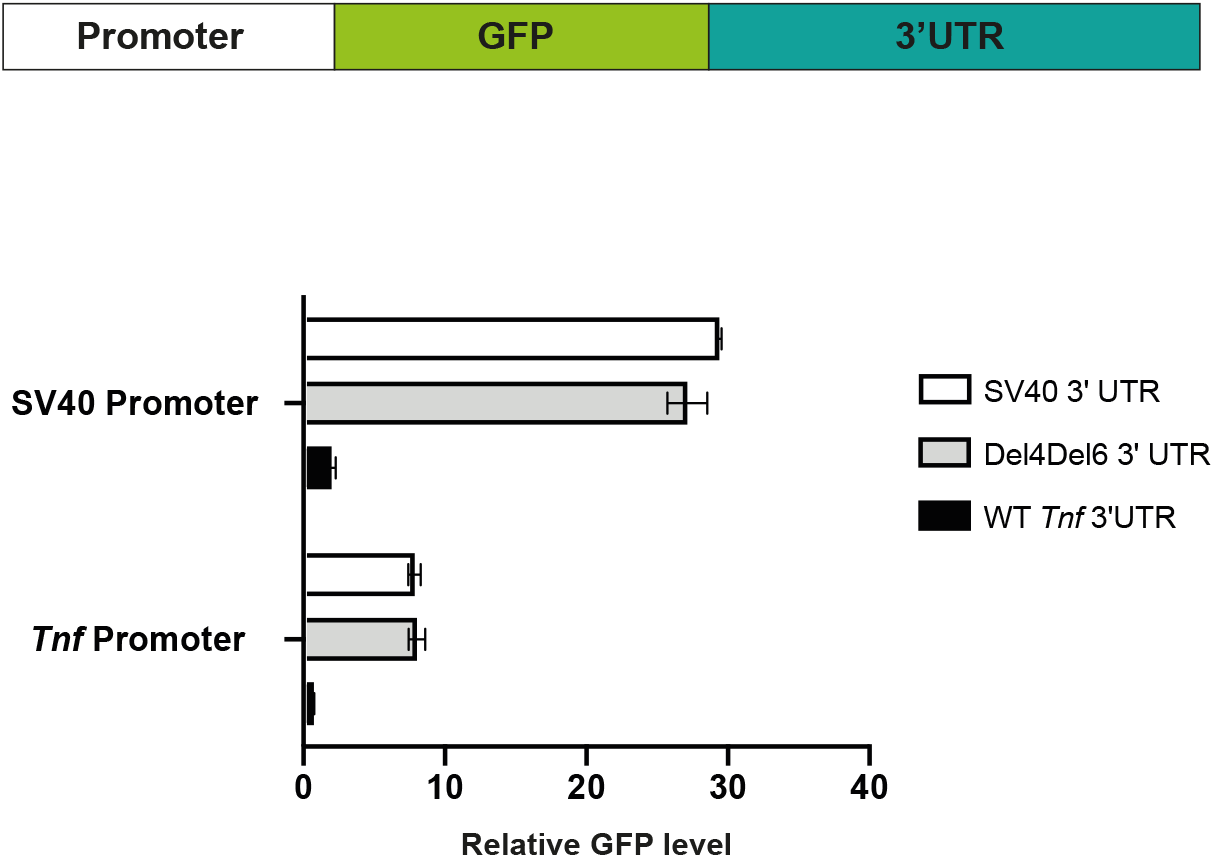
*Tnf* promoter is constitutively active in HEK293 cells. Schematic representation of the GFP reporter constructs used to assay the activity of *Tnf* promoter in HEK293 cells and relative GFP levels following transient transfection of HEK293 cells. Data represent mean ± SD from 3 independent experiments.

Video 1. **Vascular collapse in *Tnf^del4del5/del4del5^* embryos.** Label free lightsheet imaging of hemoglobin autofluorescence in cleared front paws of WT (left) and homozygote Tnf^del4del5/del4del5^ (m/m) E16.5 embryos reveals vascular fragmentation.

**Table 1.** Phenotype summary. Summary of the phenotypes observed in mice with various mutations in TNF 3’UTR.

|  | Arthritis | IBD | HVD | iBALT | BM<br>nodules | Oily<br>fur | Embryonic<br>death | Lifespan |
| --- | --- | --- | --- | --- | --- | --- | --- | --- |
| *Tnf<sup>del4/+</sup>* | - | - | - | + | + | no | no | > 1 year |
| *Tnf<sup>del4/del4</sup>* | + | +++ | - | +++ | +++ | yes | no | 300<br>days |
| *Tnf<sup>del5/+</sup>* | - | - | - | - | - | no | no | > 1 year |
| *Tnf<sup>del5/del5</sup>* | - | - | - | - | - | no | no | > 1 year |
| *Tnf<sup>del6/+</sup>* | - | - | - | - | - | no | no | > 1 year |
| *Tnf<sup>del6/del6</sup>* | - | - | - | - | - | no | no | > 1 year |
| *TNF<sup>del4del5/+</sup>* | ++ | +++ | ++ | +++ | +++ | yes | no | 200 days |
| *TNF<sup>del4del5/del4del5</sup>* | n/a | n/a | n/a | n/a | n/a | n/a | yes, E16.5 | n/a |
| *TNF<sup>del4del5/del4del5</sup>;TNFR1<sup>+/-</sup>* | +/+ | +++ | ++ | +++ | +++ | yes | no | 100 days |
| *TNF<sup>del4del6/+</sup>;TNFR1<sup>+/+</sup>* | n/a | n/a | n/a | n/a | n/a | n/a | yes, stage<br>unknown | n/a |
| *TNF<sup>del4del6/+</sup>;TNFR1<sup>+/-</sup>* | ++++ | ++ | + | + | +++ | no | 80% death<br>at birth<br>(PN1) | 20 days |
| *BPSM1<sup>m/+</sup>* | ++ | + | ++ | +++ | +++ | no | no | 200 days |
| *BPSM1<sup>m/m</sup>* | ++++ | + | ++++ | ++++ | +++ | no | no | 4-8<br>weeks |
| *TNF<sup>ΔARE/+</sup>* | ++ | ++ | ++ | ++ | + | no | no | 200 days |
| *TNF<sup>ΔARE/ΔARE</sup>* | ++++ | ++++ | ++++ | ++++ | + | no | no | 4-12<br>weeks |

